# Conditional expression of Cas9 and dCas9 in *Lucilia cuprina* reveals dCas9-associated lethality

**DOI:** 10.1101/2025.08.12.669956

**Authors:** Alexis Kriete, Tatiana Basika, Rossina Novas, Esther J. Belikoff, Maxwell J. Scott

**Author notes:** Email addresses: Alexis Kriete, Tatiana Basika, Rossina Novas, Esther J. Belikoff.

## Abstract

Conditional sex transformation systems are promising tools in the fight against insect pests. In this study, we developed and tested CRISPR-based, tetracycline-repressible sex transformation strains in the Australian sheep blowfly, *Lucilia cuprina*. Two CRISPR effector molecules, Cas9 and dCas9, were employed to target the sex-determining gene *transformer* with the goal of turning female blowflies into males. The Cas9 version of the system induced robust knockout of a visual marker gene but failed to trigger sex transformation without external provision of *transformer*-targeting sgRNAs. Furthermore, we found that dCas9 expression was linked to several deleterious phenotypes, including developmental delays, reduced body weight, and death. Our study provides the first proof-of-concept conditional CRISPR systems in *L. cuprina*, and suggests that while dCas9 is toxic at high levels in this species, Cas9 is well-tolerated and may be able to induce sex transformation with minor modifications to the system.

## Introduction

Each year, disease-vectoring insects kill over 700,000 people^1^ and agricultural insect pests cost the world’s economy an estimated $220 billion.^2^ Broad-spectrum insecticides are widely used for pest control, but can harm non-target species^3^ and lead to the rapid evolution of resistance in target species.^4,5^ Genetics-based approaches, such as the Sterile Insect Technique (SIT), can be used as more sustainable and ecologically-friendly forms of pest control.^6^ In SIT programs, insects are reared in factories, sterilized using radiation, and repeatedly released in vast quantities in a target area. Sterilized males mate with wild females, lowering the wild population’s reproductive output and leading to population suppression or elimination.^7^ Modeling^8^ and large-scale field tests^9^ have shown that SIT is more effective if only males are released, and a major focus of SIT programs has been devising methods to eliminate females from the release pool.^10,11^ One option is to use tetracycline-repressible (Tet-Off) systems to conditionally express a female-lethal gene. These systems work by using a “driver” construct to express tTA (tetracycline-controlled transactivator); tTA binds to tetO (tetracycline operator) sites upstream of an “effector” (female-lethal) gene, activating its expression.^12–14^ Tetracycline, or a related antibiotic like doxycycline, is used to turn the system off by binding to tTA and preventing effector gene expression. Highly efficient Tet-Off strains that kill females at the embryo or larval stages have been developed in several major insect pests, including the New World Screwworm (*Cochliomyia hominivorax*)^15,16^ and Australian sheep blowfly (*Lucilia cuprina*).^17,18^

An alternative to killing females would be transforming them – that is, turning XX females into XX males. Such a sex transformation system would both solve the problem of sex-sorting and effectively double the rate of male production for SIT. Modeling suggests that sex transformation systems can be more effective than female-killing systems for population suppression, provided that the XX males are sexually competitive.^19^ In *L. cuprina* and *C. hominivorax*, the *transformer* (*Lctra* and *Chtra*) gene is essential for female development, and targeted disruption of *Lctra* or *Chtra* leads to female masculinization.^20,21^ This has spurred interest in developing conditional sex transformation systems targeting *tra* in these species. RNAi lines using the Tet-Off system to knock down *Lctra* RNA were recently developed in *L. cuprina*,^22^ but fully transformed (XX) males from these lines died before adulthood, and so could not contribute to population suppression with SIT. To date, no conditional sex transformation system that produces fully-transformed adult XX males has been made in the blowfly or screwworm.

In this study, we built and tested the first conditional Cas9- and dCas9-expressing transgenic strains in *L. cuprina*, targeting *Lctra* in order to trigger female-to-male sex transformation. Cas9 disrupts its target genes by cleaving DNA at target sites and inducing loss-of-function mutations.^23^ dCas9, in contrast, knocks down gene expression by binding to the target site without cleaving it, blocking the transcriptional machinery.^24^ While Cas9 has been used in diverse insect species, dCas9 has not been tested in *L. cuprina* or more generally in other insects, and we wished to investigate its potential utility for targeted gene knockdown in blowflies as an alternative to RNAi.

We established several two-component transgenic strains by crossing Cas9 and dCas9 effector lines to tTA-expressing driver lines. RT-PCR analysis showed that all strains expressed high levels of Cas9 or dCas9 at the early embryo stage when reared off tetracycline, and we observed robust knockout of the *DsRedexpress 2 (RFPex)* fluorescent marker gene in Cas9-expressing strains. Unexpectedly, however, we found no evidence of sex transformation in any strain tested. Additional experiments strongly suggested that this was due to inefficient processing of sgRNAs in early embryos and a consequent lack of *Lctra* targeting. While high levels of Cas9 expression were well-tolerated by our flies, dCas9 expression was consistently associated with lethal or sub-lethal phenotypes, including reduced body weight and delayed development. Our research suggests that dCas9 is likely toxic to *L. cuprina* at high doses, and therefore unsuitable for use in a sex transformation system, but that a modified version of the Cas9-based system may be able to achieve efficient conditional sex transformation.

## Materials and Methods

### *In vitro* and *in vivo* assessment of sgRNAs targeting *transformer*

sgRNAs targeting *Lctra* were selected using CRISPOR (http://crispor.tefor.net/) with the *L. cuprina* ASM2204524v1 genome assembly (GenBank: GCA_022045245.1). sgRNA sequences are listed in Table S1. *In vitro* Cas9 cleavage assays were performed by combining 1 µL of 1 µM Cas9 (Alt-R™ S.p. Cas9 Nuclease V3, Integrated DNA Technologies), 3 µL of 300 nM sgRNA, and 3 µL NEBuffer™ 3.1 (New England Biolabs) in a 27 µL reaction volume. Following two 10-minute incubations at 25°C and 37°C, 50 ng template DNA was added and mixtures were incubated for 1 hour at 37°C. 1 µL of 20 µg/µL Proteinase K was added to arrest Cas9 activity before running mixtures on a gel. Two reactions, one with Cas9 and a control without Cas9, were prepared for each sgRNA. Table S2 lists primers used to amplify template DNA.

The tra3 sgRNA was further assessed *in vivo* by microinjecting wild-type (LA07 strain) *L. cuprina* embryos with a mixture of 125 ng/µL tra3 sgRNA, 250 ng/µL Cas9, 1/20th vol. NEBuffer™ 3.1, and 1/10th vol. Phenol Red dye (Sigma-Aldrich). Microinjections were performed as previously described.^25^ Intersex females were identified by inspecting injected flies for masculinization. DNA was extracted individually from intersex females by homogenizing samples in 500 µL STE buffer (10mM Tris, 1mM EDTA, 100mM NaCl, pH 8.0) in a bead mill (HT Mini: OPS Diagnostics) at 3200 RPM for 2 minutes, then purifying DNA from the homogenate using a DNEasy Blood & Tissue kit (QIAGEN) following the manufacturer’s protocol. PCR was performed on pooled DNA to amplify the region of *Lctra* encompassing the *tra3* sgRNA target site using primers listed in Table S2. The online EditCo ICE CRISPR Analysis Tool (https://ice.editco.bio/<u>#/</u>) was used to determine indel rates based on Sanger DNA sequencing data.

### Plasmid Construction

To construct the plasmids used to assess *L. cuprina U6* gene promoters, we followed a similar strategy as for the previously described *C. hominivorax U6* promoters.^26^ DNA fragments containing approximately 500bp upstream of the transcription initiation site, followed by two BbsI sites, the sgRNA sequence (targeting *Lctra*), and about 200bp of 3’ flanking from the *U6* gene were synthesized and cloned into the pUCIDT vector using Golden Gate Assembly (Table S3).

To assemble the Cas9 and dCas9 effector plasmids used for germline transformation, component fragments were first excised or amplified from source plasmids using the restriction enzymes or PCR primers listed in Table S4. The source plasmid bearing the pBac-attP-RFPex-attP-pBac fragment was derived by cloning the dsRedex2 marker from pB[Lchsp83-DsRedex2]^25^ into the pBac2 backbone.^26^ The dCas9 fragment was amplified from the source plasmid pAW91.dCas9 (Addgene: #104372).^27^ The Cas9-p10polyA and p10polyA fragments were amplified from the IattB-Nos-Cas9-P10-ZsGreen-IattB plasmid (Genbank: PV567588.1). The tetO-Hsp70 fragments originated from the pEF1 plasmid (Genbank: KT749917.1).^17^

Next, two rounds of NEBuilder® HiFi DNA Assembly were used to ligate the fragments to form the final effector plasmids, following the manufacturer’s assembly protocol. First, Cas9 or dCas9 effector plasmids lacking sgRNA arrays were assembled by ligating the following fragments: pBac-attP-RFPex-attP-pBac, Cas9-p10polyA, and tetO-Hsp70 (Cas9 effector); pBac-attP-RFPex-attP-pBac, p10polyA, dCas9, and tetO-Hsp70 (dCas9 effector). The sgRNA arrays (*tra135*, *tra246*, and *ryt*) were synthesized by Genscript and incorporated into the Cas9 and dCas9 effector plasmids by linearizing the Cas9/dCas9 plasmids with the restriction enzyme SgrDI and ligating them with PCR-amplified sgRNA array fragments. The final 6 effector plasmids will be deposited with Addgene prior to publication.

### Embryonic microinjections, germline transformation and fly rearing

The LA07 wild-type strain of *L. cuprina* was maintained as previously described.^25^ For assessment of *LccU6* promoter activity, a mix of Cas9 protein (750 ng/µL) or Cas9 plasmid (500 ng/L), U6 promoter plasmid (500 ng/mL) and Lchsp83-ZsGreen plasmid (300 ng/µL) were injected into the posterior end of preblastoderm embryos as described previously.^28^ Hatched L1 larvae that showed transient expression of ZsGreen were collected and pooled. Genomic DNA was extracted and PCR and amplicon DNA sequencing performed as previously described.^28^ The amplicon sequencing results were analyzed using CRISPResso2.^29^

For germline transformation, embryos were injected with 50 ng/µL hyperactive *piggyBac* mRNA, 350 ng/µL of Lchsp83-hyppBac helper plasmid (sequence to be deposited with Addgene prior to publication), and 250 ng/uL effector plasmid. Capped and polyadenylated hyperactive *piggyBac* RNA was synthesized using the HiScribe® T7 ARCA mRNA kit (New England Biolabs) with the Lchsp83-hyppBac plasmid as a template. Surviving G0s were crossed individually to wild-type flies, and G1 offspring were screened at the late embryo or early first instar larval stage for red fluorescence, indicating germline integration of the effector construct. Effector lines were bred to the existing driver lines DR3#6 and DR7#2^30,31^ to establish the two-component sex transformation strains tested in this study. When possible, these strains were bred to homozygosity for both the driver and effector constructs. Strains were maintained on 100 µg/mL tetracycline or 25 µg/mL doxycycline. For the experiments detailed below, flies were tested either on antibiotics (at the aforementioned doses), or off antibiotics (after removing antibiotics from the adult parents diet).

For microinjection experiments with *Lctra* sgRNAs and established Cas9- or dCas9-expressing strains, the injection mixtures containing 300mM KCl, 1/20th vol. NEBuffer™ 3.1, 1/10th vol. Phenol Red dye, and the sgRNAs tra2, tra4, and tra6 (200 ng/uL each) were injected into embryos following the earlier microinjection protocol. Flies were reared off tetracycline to ensure that Cas9 or dCas9 were expressed in early embryos. Injected embryos were reared to adulthood along with uninjected controls, and the number of males, females, and intersex flies in each group was recorded as described previously. To verify editing of *Lctra* in Cas9 strains, DNA was extracted individually from injected intersex females. DNA was also extracted from three males and females from each of the uninjected Cas9 strains and pooled in equimolar quantities to serve as controls. PCR was performed with primers flanking the sgRNA target sites in order to amplify products containing one or more large CRISPR-induced deletions. *GST-1* was amplified as a positive control. Sanger sequencing was used to verify editing of *Lctra* in injected intersex females. To determine whether injected males from the dCas9 strains were XY, DNA was extracted from each male and PCR performed using Y-linked gene primers. Table S2 lists all primers used in this experiment.

### Phenotypic assessment of sex transformation strains

To evaluate *DsRedexpress 2* (*RFPex*) knockout (in Cas9 strains) and knockdown (in dCas9 strains), third-instar (L3) larvae from strains carrying the *ryt* sgRNA array were screened for RFP intensity using a fluorescence microscope. Upon reaching adulthood, flies were inspected for sex transformation due to *Lctra* disruption (in all strains) and cuticle discoloration due to *yellow (LcY)* disruption (in strains carrying the *ryt* sgRNA array). To evaluate sex transformation, flies were scored as male (short interocular distance and male external genitalia), female (wide interocular distance and female external genitalia), or intersex (intermediate or conflicting sexual phenotype, e.g. masculinized genitalia with female interocular distance). As no intersex flies were ever observed, the adult sex ratio was defined as the proportion of males. The eclosion rate, defined as the proportion of pupae surviving to adulthood, was also recorded for each replicate. To evaluate knockout/knockdown of *yellow*, adults were inspected for characteristic brown wings and body color.^21^ Differences in the mean adult sex ratio and eclosion rate across different experimental groups were statistically analyzed in R (v. 4.4.1) using pairwise t-tests with Benjamini-Hochberg correction for multiple comparisons. The raw data from sex ratio and eclosion rate experiments is provided in Table S5.

Additional experiments to measure larval development rates and pupal weight were performed with a subset of sex transformation strains. For each strain, independent colonies containing approximately 120 first-instar larvae were set and reared to adulthood on or off 100 µg/mL tetracycline (N=3 replicate colonies per condition). To assess delays in larval development, the number of wandering-stage third instar (L3) larvae that emerged from the larval meat diet each day was recorded cumulatively until the last larva had emerged. Larval counts were then converted to proportions (as raw counts could not be directly compared statistically, due to small natural variations in colony size) and charted to visualize larval developmental trajectories. For the pupal weight experiments, up to 50 pupae from each replicate colony were weighed, and the average pupal weight was calculated from these values and plotted. A few strains yielded fewer than 50 pupae per replicate due to low survival rates; this information is reported in Table S6. Statistical comparisons of mean pupal weights across experimental groups were performed in R (v. 4.4.1) using pairwise t-tests with Benjamini-Hochberg correction for multiple comparisons.

### RNA extraction and RT-PCR

RNA was extracted from pooled 2-6 h embryos and 24 h (first instar) larvae from sex transformation strains reared on or off tetracycline. To extract RNA, samples (N ≥ 20 individuals per sample) were placed in Trizol™ (Thermo-Fisher) and homogenized in a bead mill for two 60s cycles at 3200 RPM. RNA was isolated from the homogenized samples following the QIAGEN RNeasy Mini Kit protocol for total RNA extraction from animal tissue. cDNA was synthesized using the SuperScript™ III First-Strand Synthesis SuperMix (Invitrogen) protocol. A control reaction lacking the reverse transcriptase enzyme (SuperScript III) was run alongside each RNA sample. RT-PCR was performed using primer pairs listed in Table S2.

### Evaluation of indel rates at target sites in Cas9 strains

DNA was isolated (as previously described) from 2-6 h old embryos (N≥20 individuals per sample), 24 h old first instar larvae (N≥10 individuals per sample), and 2 d old adult flies (N≥5 individuals per sample). PCR was used to amplify regions flanking the target sites of the following sgRNAs: RFPex, yellow, tra2, tra4, tra6, and tra7. PCR primers are listed in Table S2. Amplicon sequencing and indel rate determination was performed by Azenta Life Sciences.

## Results

### Design of conditional sex transformation systems in *L. cuprina*

We designed six effector constructs (Fig. 1A) carrying Cas9 or dCas9 under the control of the *tetO*_21_-*Lchsp70* enhancer-promoter used previously.^25^ In the absence of tetracycline, tTA binds to tetO sites in the effector and induces expression of Cas9 or dCas9. Effector constructs also carried a constitutively expressed red fluorescent marker gene, *DsRedexpress 2* (*RFPe*x), driven by the *Lchsp83* gene promoter. A tRNA-based processing strategy^32^ was used to express multiple sgRNAs from a single promoter. To identify a suitable RNA polymerase III promoter, we used a transient expression assay to assess the promoters from three *U6* genes identified in the *L. cuprina* genome.^33^ The assay was similar to that used previously for *C. hominivorax* U6 promoters^28^ except that the U6 plasmids expressed a sgRNA targeting *Lctra*. The results indicated the *LccU6b* promoter was the most active in *L. cuprina* embryos (Table S7). Each construct contained one of three sgRNA arrays (*tra135*, *tra246*, or *ryt*), consisting of three different sgRNAs (spacer-scaffold sequences) separated by tRNAs driven by the *LccU6b* promoter. Endogenous cellular machinery cleaves tRNAs from the precursor transcript, liberating individual sgRNAs.^32^

**FIG 1.**
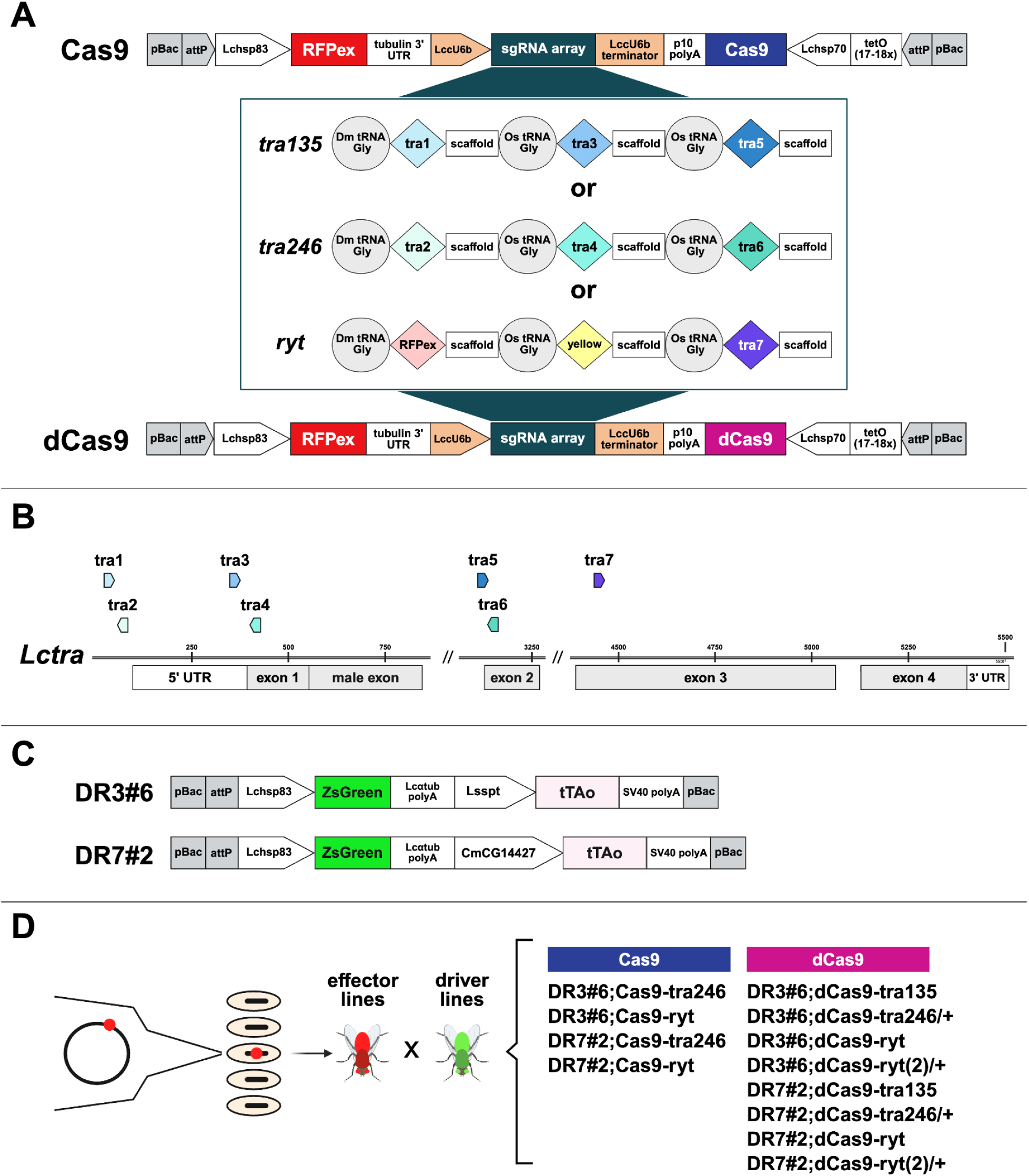
Design of conditional sex transformation systems in *L. cuprina*. **(A)** Schematic illustrations of Cas9- and dCas9-bearing effector constructs. Components of the three sgRNA arrays (*tra135*, *tra246*, and *ryt*) are shown in the middle panel. Expression of Cas9 or dCas9 is controlled by the *tetO-Lchsp70* enhancer-promoter. The constitutive promoter *Lchsp83* drives expression of the red fluorescent marker DsRedexpress 2 (*RFPex*). The *U6* (RNA Polymerase III) promoter *LccU6b* drives expression of the sgRNA arrays. **(B)** Map of the *L. cuprina transformer* gene (*Lctra*) showing the locations of target sites for sgRNAs tra1 - tra7. Arrow orientation corresponds to the DNA strand targeted by the Cas9/dCas9-sgRNA complex (rightward-facing arrow: targets non-coding strand; leftward-facing arrow: targets coding strand). **(C)** Schematic illustrations of *tTAo*-expressing constructs in the driver lines DR3#6 and DR7#2. Expression of *tTAo* is controlled by the *Lsspt* promoter in DR3#6 and by the *CmCG14427* promoter in DR7#2. Expression of the green fluorescent marker *ZsGreen* is controlled by the constitutive promoter *Lchsp83*. **(D)** Overview of sex transformation strains established and tested in this study. Effector lines (red) were created using *piggyBac*-mediated transgenesis and crossed to driver lines (green). The names of the resulting two-component (driver + effector) sex transformation strains are shown on the right.

The three sgRNAs encoded by the *ryt* sgRNA array (*RFPex*, *yellow*, and *tra7*) were chosen to validate our system because they were previously shown to efficiently disrupt their target genes and produce visible knockout phenotypes in *L. cuprina*.^21,22,28^ The *RFPex* and *yellow* sgRNAs target the *RFPex* and *yellow* (*LcY)* genes, while the *tra7* sgRNA targets the third exon of *Lctra* (Fig. 1B). The sgRNA arrays *tra135* and *tra246* exclusively expressed sgRNA targeting *Lctra* (Fig. 1B). The sgRNAs *tra1* and *tra2* were designed to target the promoter, because dCas9-mediated transcriptional repression may be more efficient if dCas9 is directed to this region.^34^ The sgRNAs *tra2*, *tra4*, and *tra6* target the coding strand, which may also enhance dCas9’s performance.^34^ sgRNAs were evaluated using Cas9 cleavage assays, which revealed that every sgRNA was capable of efficiently cleaving its target *in vitro* (Fig. S1A). One of the sgRNAs, *tra3*, was further validated *in vivo* by coinjecting embryos with Cas9 and *tra3* sgRNA, resulting in high (85%) editing rates at the target site in injected intersex females (Fig. S1B).

The driver constructs DR3#6 and DR7#2 (Fig. 1C) were used to express tTA under the control of the promoters *Lsspt* (DR3) and *CmCG14427* (DR7), which are highly active in early embryos.^30,31^ The DR3 driver lines also shows significant expression at later developmental stages and in adult ovaries.^30,31^ Both drivers carry a constitutively-expressed green fluorescent marker (*ZsGreen*) driven by the *Lchsp83* gene promoter.

### Establishment of transgenic lines

Effector plasmids were injected into wild-type *L. cuprina* embryos alongside hyperactive *piggyBac* transposase mRNA and DNA helper plasmid. We used the hyperactive version of *piggyBac* because it has been linked to much higher germline transformation rates in insects than conventional *piggyBac*.^35^ However, in our hands, transformation rates with hyperactive *piggyBac* were low (∼4%). We obtained two Cas9 lines (Cas9-ryt and Cas9-tra246) and four dCas9 lines (dCas9-tra135, dCas9-tra246, and two independent insertions of the dCas9-ryt effector, dubbed dCas9-ryt and dCas9-ryt(2)) from *piggyBac* injections. These six effector lines were crossed to the driver lines DR3#6 and DR7#2 to establish the final set of 12 two-component (driver/effector) sex transformation strains tested in this study (Fig. 1D).

Two-component sex transformation strains were maintained on 100 µg/mL tetracycline or 25 µg/mL doxycycline to repress Cas9/dCas9 expression. However, flies homozygous for the dCas9-tra246 or dCas9-ryt(2) effector constructs died *en masse* during the embryo or early larval stages, even when reared on antibiotics, presumably due to leaky dCas9 expression and associated toxicity. These strains could only be maintained in the hemizygous condition (i.e. homozygous for the driver, heterozygous for the effector). To test whether these strains could be made fully homozygous on a higher dose of antibiotics, we reared a subset of hemizygous flies on 100 µg/mL doxycycline (4x the standard dose) and screened their offspring for dCas9 homozygotes bearing two copies of the *RFPex* marker. However, we observed nearly complete lethality among larvae homozygous for dCas9 in three of the four strains tested (Table S8).

### RT-PCR analysis of gene expression in sex transformation strains

We performed RT-PCR to evaluate expression of *Cas9* and *dCas9* in sex transformation strains reared on and off tetracycline. Both genes were robustly expressed in 2-6 h old embryos and 24 h larvae reared off tetracycline (Fig. 2A, Fig. S2A-F). Minimal leaky expression of *Cas9* or *dCas9* was observed in most strains reared on tetracycline, with the exception of the DR3#6;dCas9-ryt(2)/+ and DR7#2;dCas9-ryt(2)/+ strains, in which tetracycline appeared to be unable to repress *dCas9* expression (Fig. S2F). To determine whether Cas9 and dCas9 were altering the course of sexual development by disrupting *Lctra* expression, we also performed RT-PCR using primers designed to amplify sex-specifically spliced *Lctra* transcripts. As *Lctra* splicing is auto-regulated, disruption of *Lctra* with Cas9 or dCas9 should lead to increased expression of the male *Lctra* transcript overall.^20^ However, we did not observe any noticeable shifts in male vs. female *Lctra* transcript expression patterns. Similarly, we used RT-PCR to evaluate *dCas9*-mediated knockdown of the target genes *RFPex* and *yellow (LcY)*, but observed no change in the expression of these genes in flies reared off tetracycline. These results suggested that although *Cas9* and *dCas9* could be expressed at high levels early in development using our system, these effector molecules were not efficiently disrupting their target genes. This may have been caused by inadequate sgRNA expression. To determine if sgRNA arrays were being transcribed early in development, we performed RT-PCR using primers targeting the sgRNA tracer and scaffold regions (Fig. 2B). These primers can use either the uncleaved, precursor sgRNA array transcript as a template, or individual, cleaved sgRNAs. In 2-6 h old embryos, we observed amplification of longer products which could only have come from the unprocessed sgRNA array precursor transcript. By 24h, only short products were amplified, which may indicate that sgRNA arrays are more efficiently processed at this stage. Regardless, we confirmed that the *LccU6b* promoter was actively expressing sgRNA arrays in early embryos and larvae.

**FIG 2.**
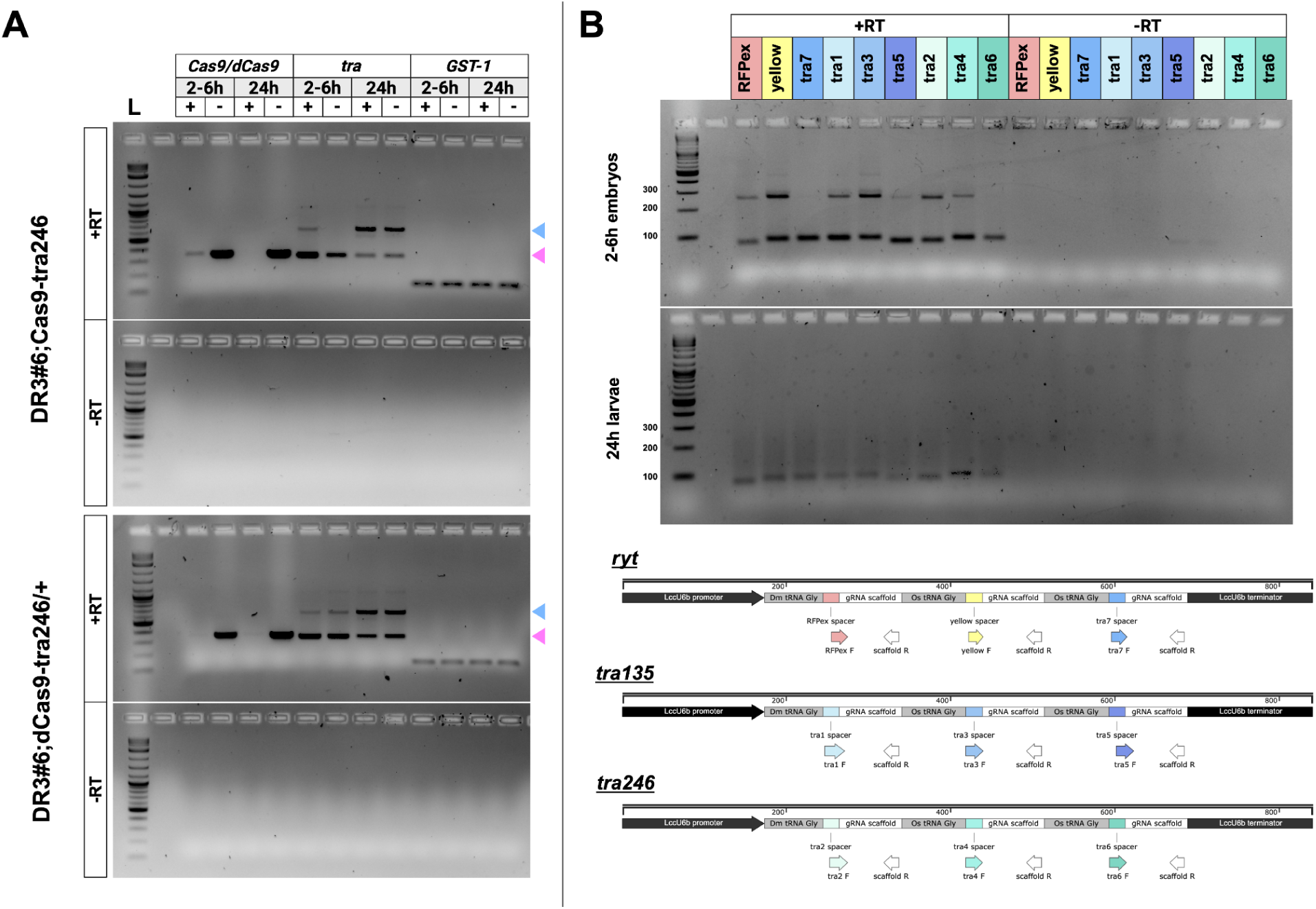
Expression of Cas9/dCas9, sgRNA array transcript, and target genes in embryos and first-instar larvae from sex transformation strains. (A) RT-PCR analysis of Cas9/dCas9 and Lctra expression patterns in 2-6 h old embryos and first-instar larvae from representative Cas9/dCas9-expressing sex transformation strains. Gel images from additional strains are displayed in Figure S2. Blue and pink arrows indicate expected product sizes of male and female splice variants of Lctra, respectively. The GST-1 primer pair is used as a positive control. Lanes labeled with + or - denote samples from flies reared on or off 100 µg/mL tetracycline, respectively. +RT = reactions run with reverse transcriptase; -RT = reactions run without reverse transcriptase (negative controls). L = DNA ladder. (B) RT-PCR analysis of sgRNA array and/or individual sgRNA expression patterns in 2-6 h old embryos (top row) and first-instar larvae (bottom row) from sex transformation strains. Maps of the sgRNA arrays and RT-PCR primer binding locations are shown below the gel. Larger (∼290 bp) products can only be amplified using the uncleaved sgRNA array primary transcript as a template, whereas shorter (∼90 bp) products can be amplified from either the uncleaved sgRNA array primary transcript or from individual sgRNAs.

### Conditional knockout of *RFPex* in Cas9-expressing strains

Fly strains carrying the *ryt* sgRNA array should express sgRNA targeting the fluorescent marker *RFPex*. To assess the effectiveness of knockout (*Cas9*) and knockdown (*dCas9*) of *RFPex* using our conditional gene expression system, we screened third instar (L3) larvae from strains reared on and off tetracycline or doxycycline (Fig 3). We observed consistent reductions in red fluorescence in larvae from Cas9 strains reared off antibiotics, regardless of the driver or antibiotic tested. Conversely, larvae from dCas9 strains reared off antibiotics did not display any noticeable reduction in RFP intensity relative to antibiotic-reared controls.

**FIG 3.**
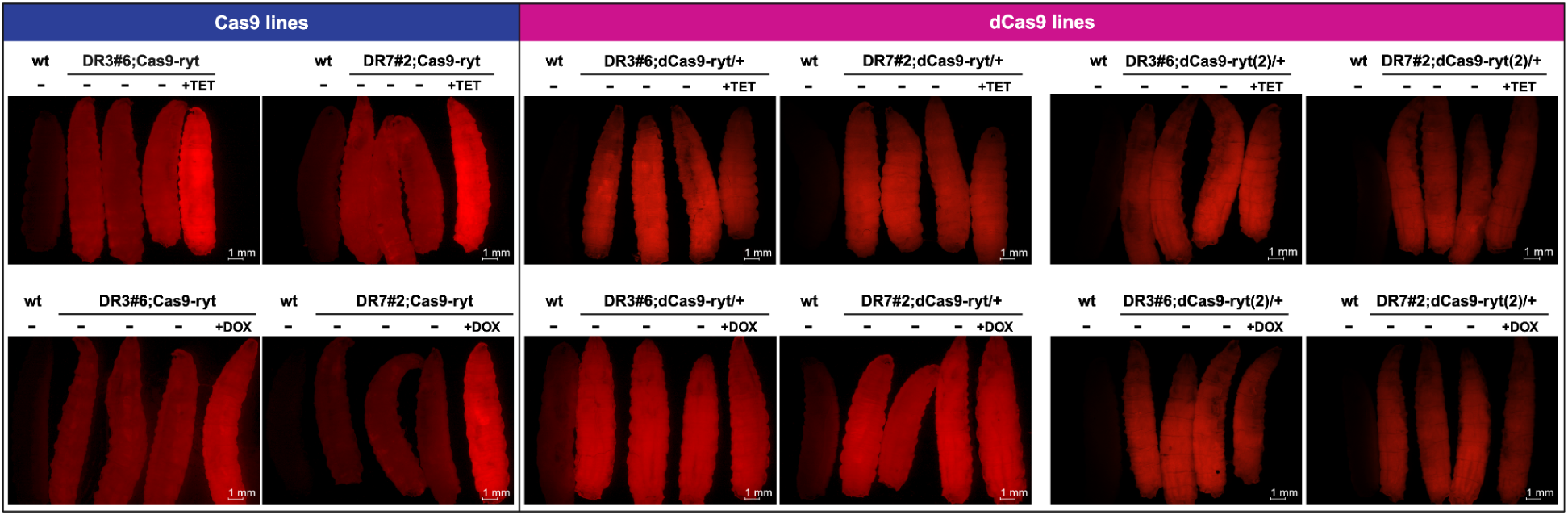
Conditional expression of Cas9 leads to knockout of RFPex and loss of red fluorescence in third-instar larvae. “+TET” and “+DOX” indicate flies reared on 100 µg/mL tetracycline or 25 ug/mL doxycycline, respectively, while “-” labels denote flies reared off antibiotics. “wt” = wild-type (non-fluorescent) larva.

### Phenotypic evaluation of sex transformation and *yellow* disruption

Flies were reared to adulthood to be screened for disruption of the target genes *Lctra* and *yellow* (*LcY*). Pupal eclosion rates were recorded prior to screening. Unexpectedly, we found that all four of the homozygous dCas9 strains tested displayed complete or partial lethality at the pupal stage when reared off antibiotics (Fig. 4A). This dCas9-associated lethality appeared to be dose-dependent, as the hemizygous dCas9 strains displayed no or smaller reductions in the eclosion rate when reared off antibiotics. Lethality rates were higher in strains reared off tetracycline than off doxycycline, presumably because doxycycline is a more stable molecule^36^ and may have been maternally inherited. In contrast to the lethality observed in dCas9 strains, the four homozygous Cas9 strains had universally high (83-97%) eclosion rates whether on and off antibiotics (Fig. 4A), suggesting that pupal lethality was due to dCas9 expression specifically.

**FIG 4.**
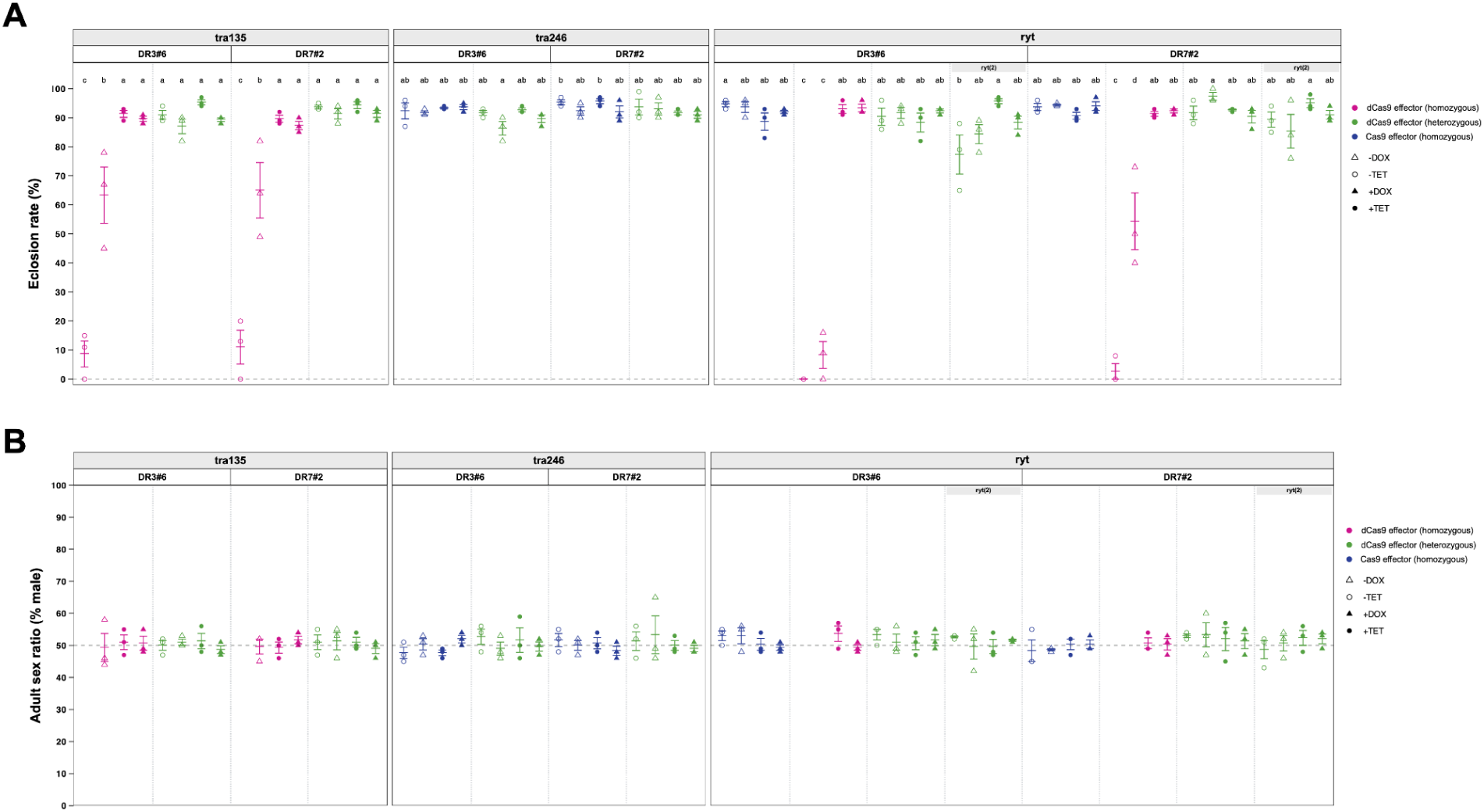
Eclosion rates and adult sex ratios of sex transformation strains reared on or off antibiotics. **(A)** Eclosion rates (percentage of pupae surviving to adulthood). **(B)** Adult sex ratios (percentage male). Labels above the plots indicate the sgRNA array (*tra135*, *tra246*, or *ryt*) and driver (DR3#6 or DR7#2) in each sex transformation strains tested. Data points are color-coded according to the identity and zygosity of the Cas effector (pink = homozygous for d*Cas9*; green = heterozygous for *dCas9*; blue = homozygous for *Cas9*). Filled circles and triangles represent strains reared on 100 µg/mL tetracycline or 25 µg/mL doxycycline (respectively); unfilled circles and triangles represent strains reared off tetracycline or doxycycline. Horizontal lines with error bars indicate the mean of the replicates ± SE for each experimental group. Each replicate consisted of N ≥ 50 flies, with the exception of the following strains: DR3#6;dCas9-tra135 (-TET); DR7#2;dCas9-tra135 (-TET), DR3#6;dCas9-ryt (-TET and -DOX); DR7#2;dCas9-ryt (-TET and -DOX). Fewer than 15 flies per replicate survived to adulthood in these groups, so sex ratios were not calculated and displayed on the plot, but the raw counts can be found in Table S4. Statistically significant differences in eclosion rates across experimental groups are indicated by different letter combinations (p < 0.05; unpaired t-test with Benjamini-Hochberg correction for multiple comparisons).

Surviving adults were screened for sex transformation phenotypes. CRISPR-mediated knockout or knockdown of *Lctra* should cause partial or complete masculinization, leading to visibly intersex females and/or a male-biased sex ratio in strains reared off antibiotics. Unexpectedly, no intersex females were ever observed, and no strain deviated strongly from the baseline 50:50 sex ratio (Fig. 4B). We also screened flies from strains carrying the *ryt* sgRNA array, which should produce sgRNA targeting *yellow*, for brown wing and cuticle discoloration indicative of biallelic disruption of the *yellow* gene.^21^ As with *Lctra*, we did not observe any adults displaying a brown body mutant phenotype in our screens.

### dCas9 expression is associated with delayed larval development and reduced body mass

While raising flies for phenotypic screening, we noticed that individuals from homozygous dCas9 strains reared off antibiotics were smaller and slower to develop than antibiotic-reared controls. To investigate these phenotypes more rigorously, we conducted follow-up experiments with a subset of sex transformation strains reared on or off 100 µg/mL tetracycline. We found that flies homozygous for dCas9 took several days longer on average to reach the third instar larval stage when reared off tetracycline compared to tetracycline-reared controls (Fig. 5A). In contrast, developmental times for hemizygous dCas9 and homozygous Cas9 strains were similar whether reared on or off tetracycline. To determine whether dCas9 expression affected body mass, weight of pupae from sex transformation strains reared on or off tetracycline was measured. We found that pupae from homozygous dCas9 strains reared off tetracycline were significantly (∼40%) lighter on average than pupae from all other groups (Fig. 5B). These data, combined with the high pupal lethality rates observed previously, strongly suggest that dCas9 is toxic to *L. cuprina* when expressed at high levels, resulting in severe developmental impairments or death.

**FIG 5.**
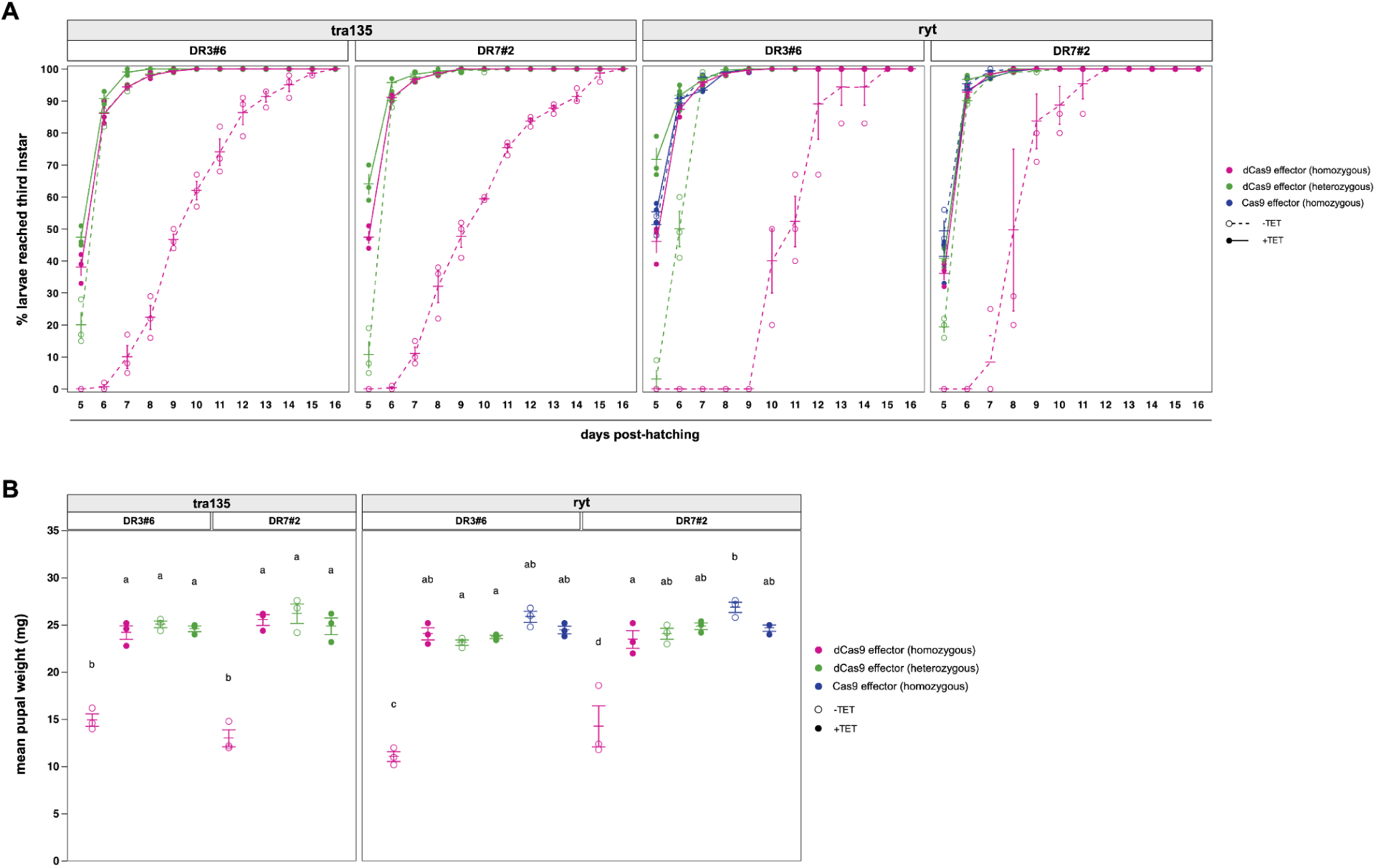
dCas9 expression is associated with delayed larval development and reduced pupal weight. **(A)** Time course showing the cumulative percentage of larvae from different sex transformation strains progressing to the third instar stage. Filled circles connected by solid lines represent flies reared on 100 µg/mL tetracycline, while unfilled circles connected by dashed lines represent flies reared off tetracycline. Horizontal lines with error bars indicate the mean of the replicates ± SE for each experimental group. 3 replicate colonies were assessed for each group. Labels above the plots indicate the sgRNA array (*tra135* or *ryt*) and driver (DR3#6 or DR7#2) in each sex transformation strains tested. Data points and lines are color-coded according to the identity and zygosity of the Cas9 effector (pink = homozygous for d*Cas9*; green = heterozygous for *dCas9*; blue = homozygous for *Cas9*). **(B)** Mean weight of pupae from different sex transformation strains reared on or off 100 µg/mL tetracycline. 3 replicate colonies were assayed for each experimental group. N=50 pupae were used per replicate for all but 3 strains, in which pupation rates were lower than average (see Table S5 for sample sizes). Horizontal lines with error bars indicate the mean of the replicates ± SE for each group. Statistically significant differences between experimental groups are indicated by different letter combinations (p < 0.05; unpaired t-test with Benjamini-Hochberg correction for multiple comparisons).

### Evaluation of indel rates in Cas9 strains

We were puzzled by the apparent failure of Cas9 or dCas9 to disrupt the *Lctra* and *yellow* genes in our study, given that Cas9 could reliably knock out *RFPex*. We noted that the *RFPex*-targeting sgRNA was in the first position of its sgRNA array, upstream of sgRNAs targeting *yellow* and *Lctra* (Fig. 1A). This raised the possibility that an sgRNA’s effectiveness might be influenced by its position in the array. Consequently, DNA was extracted from Cas9 strains reared on or off tetracycline and indel rates measured at different sgRNA target sites (Fig. 6A-B, Table S9). Indel rates were generally low (<10%) in embryos and first-instar larvae but higher in adults, implying that Cas9 persisted into adulthood and/or that sgRNA expression increased over time. Indel rates were negligible in samples reared on tetracycline, consistent with our RT-PCR data showing minimal leaky expression of *Cas9*. In all samples, indel rates at sites targeted by the first sgRNA in an array were noticeably higher than sites targeted by the second or third sgRNA.

**FIG 6.**
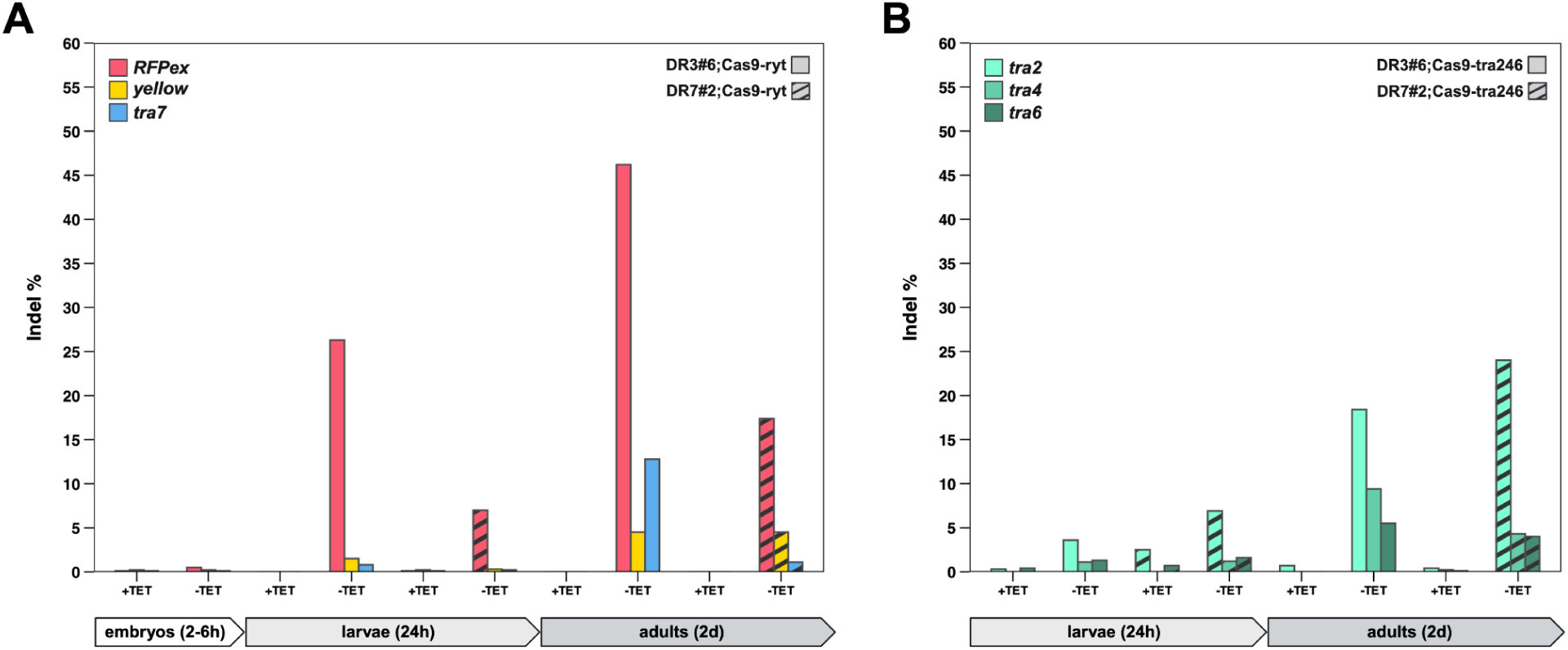
Indel rates at sgRNA target sites in Cas9-expressing strains. Indel rates at (**A**) *RFPex*, *yellow*, and *tra7* sgRNA target sites in DR3#6;Cas9-ryt and DR7#2;Cas9-ryt strains and (**B**) *tra2*, *tra4*, and *tra6* sgRNA target sites in DR3#6;Cas9-tra246 and DR7#2;Cas9-tra246 strains. Flies were sampled at three different developmental stages: 2-6 h old embryos, 24 h (first-instar) larvae, and 2d old adults. +TET and -TET indicate fly strains reared on or off 100 µg/mL tetracycline, respectively. Indel rates were calculated from amplicon sequences from pooled samples of N ≥ 5 flies/sample.

### Injection of sgRNAs targeting *tra* masculinizes Cas9-expressing females

To confirm that the lack of sex transformation in Cas9-expressing flies was due to inadequate sgRNA expression in early embryos, we injected embryos from the DR3#6;Cas9-tra246 and DR7#2;Cas9-tra246 strains with a cocktail of *Lctra*-targeting sgRNAs (200 ng/uL each of tra2, tra4, and tra6). We also injected embryos from the corresponding hemizygous dCas9 strains (DR3#6;dCas9-tra246/+ and DR7#2;dCas9-tra246/+) with the same sgRNA cocktail to determine whether dCas9 could knock down *Lctra* if flies were provided with external sgRNAs. 16% and 38% of injected females from the DR3#6;Cas9-tra246 and DR7#2;Cas9-tra246 strains, respectively, were identified as intersex based on evaluation of external genitalia (Fig. 7A). No intersex females were observed in injected dCas9 strains or in uninjected controls from the Cas9 or dCas9 strains (Table S10). Sex ratios in the injected dCas9 strains appeared heavily male-biased (Table S10), suggesting possible female-to-male transformation due to *Lctra* knockdown, but molecular genotyping using Y-linked primers confirmed that all phenotypically male flies were XY (Fig. S3). PCR amplification and Sanger sequencing of *Lctra* amplicons containing large CRISPR-induced deletions confirmed that Cas9 had successfully edited *Lctra* in intersex females (Fig. 7B).

**FIG 7.**
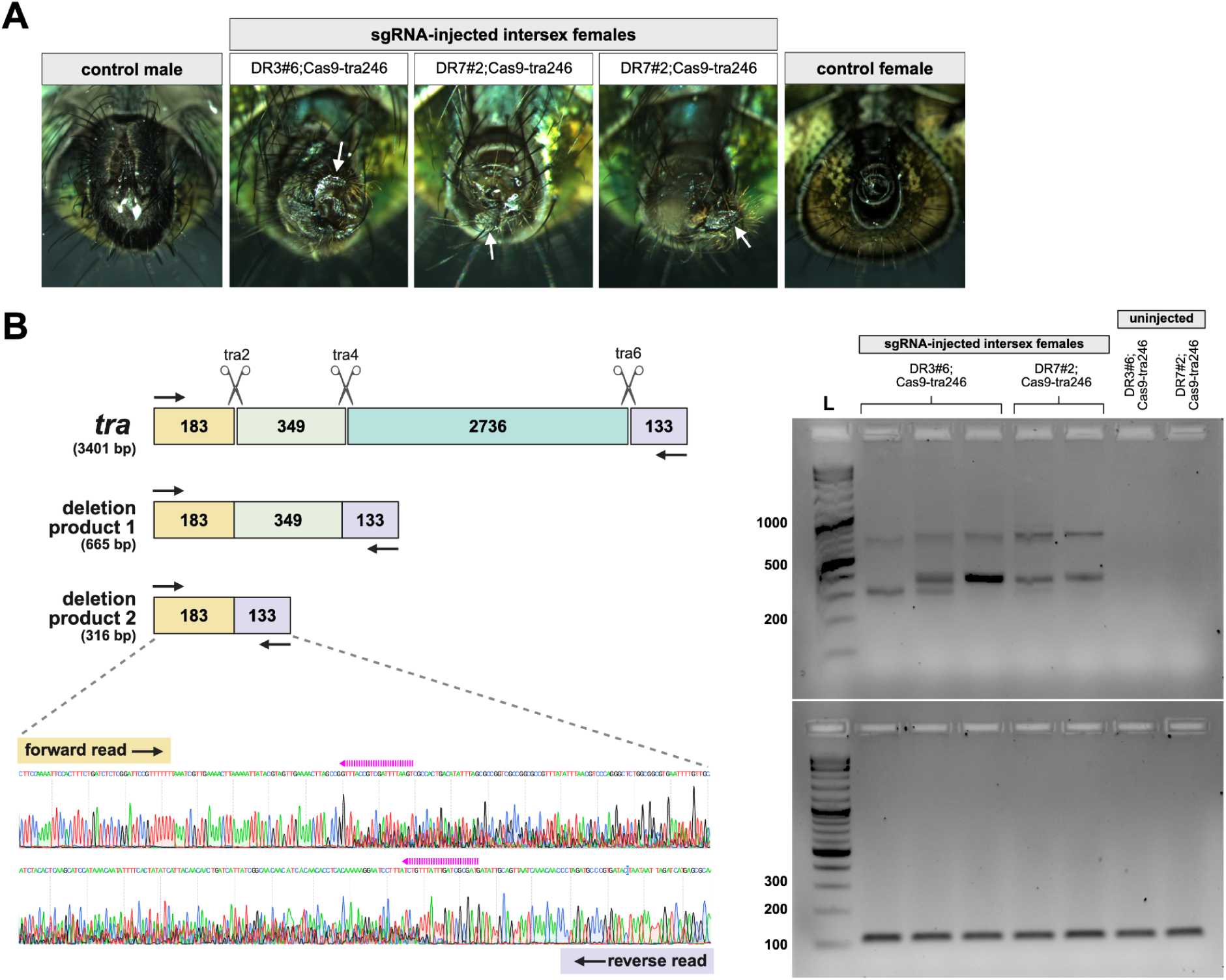
Masculinization of females from Cas9-expressing strains injected with sgRNAs targeting *Lctra.* **(A)** Ventral view of external genitalia from uninjected wild-type male (left), sgRNA-injected intersex females (middle images), and uninjected wild-type female (right). White arrows indicate malformed ovipositors in intersex females. **(B)** Molecular confirmation of Cas9-mediated editing of target sites in *Lctra* in sgRNA-injected intersex females. Left: Schematic illustration of the *Lctra* gene with Cas9 cut sites for sgRNAs *tra2*, *tra4*, and *tra6* (scissor icons). Numbers indicate the size (in base pairs) of intervening fragments. Black arrows indicate primers used for PCR, which amplify short deletion products only if Cas9-mediated cleavage has occurred at two or more cut sites. Bottom left: Representative chromatograms from Sanger sequencing of short (316 bp) deletion product from injected intersex DR3#6;Cas9-tra246 female using indicated forward and reverse primers. Dashed pink arrows above the chromatograms indicate target sites for sgRNAs *tra2* (forward read) and *tra6* (reverse read). Right: Gels showing products of PCR of *Lctra* using sgRNA target site-flanking primers (top) and *GST-1* primers as positive controls (bottom). L = DNA ladder.

## Discussion

Conditional sex transformation strains have the potential to improve SIT and other genetics-based pest control methods. In this study, we report the first conditional expression system for Cas9/dCas9 in a pest insect. More specifically, we lay the groundwork for the development of CRISPR-based sex transformation systems in the blowfly *L. cuprina*.

Successful gene disruption and sex transformation using our system depends on the robust early expression of Cas9 or dCas9 in flies reared off tetracycline, as well as the efficient expression and processing of sgRNA arrays to liberate individual sgRNAs capable of complexing with the Cas effector protein. Cas9 and dCas9 were indeed expressed at high levels in early embryos using the Lchsp70-tetO enhancer-promoter, as were sgRNA arrays using the constitutive *LccU6b* promoter. Cas9/dCas9 expression was efficiently repressed in most strains through the addition of tetracycline to the diet, a key requirement for maintaining conditional sex transformation strains in an SIT program. One target gene, the fluorescent marker *RFPex*, was reliably disrupted in all four Cas9 strains tested off tetracycline. However, despite screening thousands of flies, we never observed disruption of the other two target genes, *Lctra* and *yellow*. One possible explanation is that sgRNAs targeting those genes are less efficient than the sgRNA targeting *RFPex*. However, *in vivo* studies with *L. cuprina* indicate that the sgRNAs have comparable CRISPR editing rates (RFPex: 78%;^28^ *yellow*: 68%;^21^ *Lctra* (tra3): 85% (Fig. S1)). Furthermore, the *tra7* sgRNA has previously been reported to reliably masculinize females,^22^ and our injection experiments showed that externally-provided *Lctra*-targeting sgRNAs were sufficient to induce masculinization in Cas9-expressing flies reared off tetracycline. We therefore suspect that the lack of sex transformation observed in our strains stemmed from inefficient processing of the sgRNA arrays. We found that sgRNAs in the second or third position of the sgRNA array (relative to the promoter) were highly inefficient (0-13% editing efficiency for all conditions tested), whereas editing efficiency at sites targeted by sgRNAs in the first position of the array were always higher. The sgRNA targeting *RFPex* occupies the first position of its sgRNA array, which may explain why we saw consistent *RFPex* knockout phenotypes in flies reared off antibiotics. On the other hand, the sgRNAs *tra1* and *tra2*, which occupy the first positions of the *tra135* and *tra246* arrays respectively, target the 5’ UTR (*tra1*) or promoter (*tra2*) of *Lctra* instead of its coding sequence. Mutation of these upstream elements alone would likely not result in masculinization. We used sgRNA arrays because multiplexed sgRNA expression is highly efficient in *Drosophila*^32^ and has the benefit of reducing construct size, which is negatively correlated with germline transformation rates with *piggyBac*.^37^ However, our results suggest that CRISPR efficiency may be improved in *L. cuprina* if sgRNAs are expressed from individual promoters.

While we observed consistent *RFPex* knockout in Cas9 strains, we found no visible or molecular evidence of knockdown of any target gene in the dCas9 strains. Although we could not directly compare Cas9 and dCas9 because the effector constructs’ different genomic positions may have influenced their expression, our results suggest that dCas9 is a less potent effector than Cas9 in *L. cuprina*. Fusing dCas9 to a protein domain that inhibits gene expression in *L. cuprina* as is commonly done to control expression in human cells^38^ could potentially provide more effective inhibition of *Lctra* expression.

Our assessment of the dCas9 strains was complicated by the fact that flies homozygous for the dCas9 effector often died before adulthood. Of the eight dCas9;tTA two-component strains evaluated in this study, four could not be maintained as double-homozygous driver/effector crosses on the standard dose of tetracycline due to near-complete lethality at the embryo or larva stage. However, some homozygous dCas9 strains survived when reared on high rates of doxycycline (Table S8), suggesting that lethality resulted from high levels of dCas9 expression, rather than from the effector construct *per se*. Other dCas9;tTA strains could be maintained as double homozygotes on standard antibiotic doses without apparent ill effects, but not when tetracycline was omitted from the diet. The strains displayed severe, deleterious phenotypes – ranging from reduced pupal weight to delayed development to 100% lethality at the pupal stage – when reared off antibiotics. None of these phenotypes were observed in any of the homozygous Cas9 strains. Cas9 and dCas9 overexpression are known to be toxic in *D. melanogaster*, even in the absence of gRNAs^39,40^. However, in our *L. cuprina* strains, Cas9 appeared to be much more well-tolerated than dCas9. This is difficult to explain, given that the two effectors are nearly identical, with dCas9 differing from Cas9 only by two amino acid substitutions.^24^ However, dCas9, unlike Cas9, never cleaves its target site and destroys its own recognition mechanism, so dCas9 may repeatedly bind to DNA, leading to greater rates of replication arrest and associated lethality than Cas9. Support for this hypothesis comes from evidence in prokaryotic and eukaryotic *in vitro* DNA replication models, where dCas9 was shown to efficiently block DNA synthesis by stalling replication forks, and to bind target DNA for an estimated >44 hours.^41^ Numerous studies have shown that stalled replication forks lead to replication stress and associated genomic instability, DNA damage, and cell death.^42–44^ While it is beyond the scope of this study to investigate possible mechanisms of dCas9-associated lethality, based on our results, researchers considering the use of dCas9 expression systems in insects may wish to exercise caution.

Overall, our data suggest that minor modifications to the Cas9-based version of this system may result in improved sex transformation rates, potentially creating a powerful tool to help control blowflies and other major insect pests.

## Authorship confirmation/contribution statement

**Alexis Kriete:** data curation, formal analysis, funding acquisition, investigation, methodology, visualization, writing-original draft preparation. **Tatiana Basika:** investigation, methodology, data curation, validation, formal analysis, writing-review & editing. **Rosina Novas:** investigation, methodology, data curation, validation, formal analysis, writing-review & editing. **Esther J. Belikoff:** investigation, methodology. **Maxwell Scott**: conceptualization, funding acquisition, project administration, supervision, writing-review & editing.

