## Supplementary Figures for "Conditional expression of Cas9 and dCas9 in *Lucilia cuprina* reveals dCas9-associated lethality"

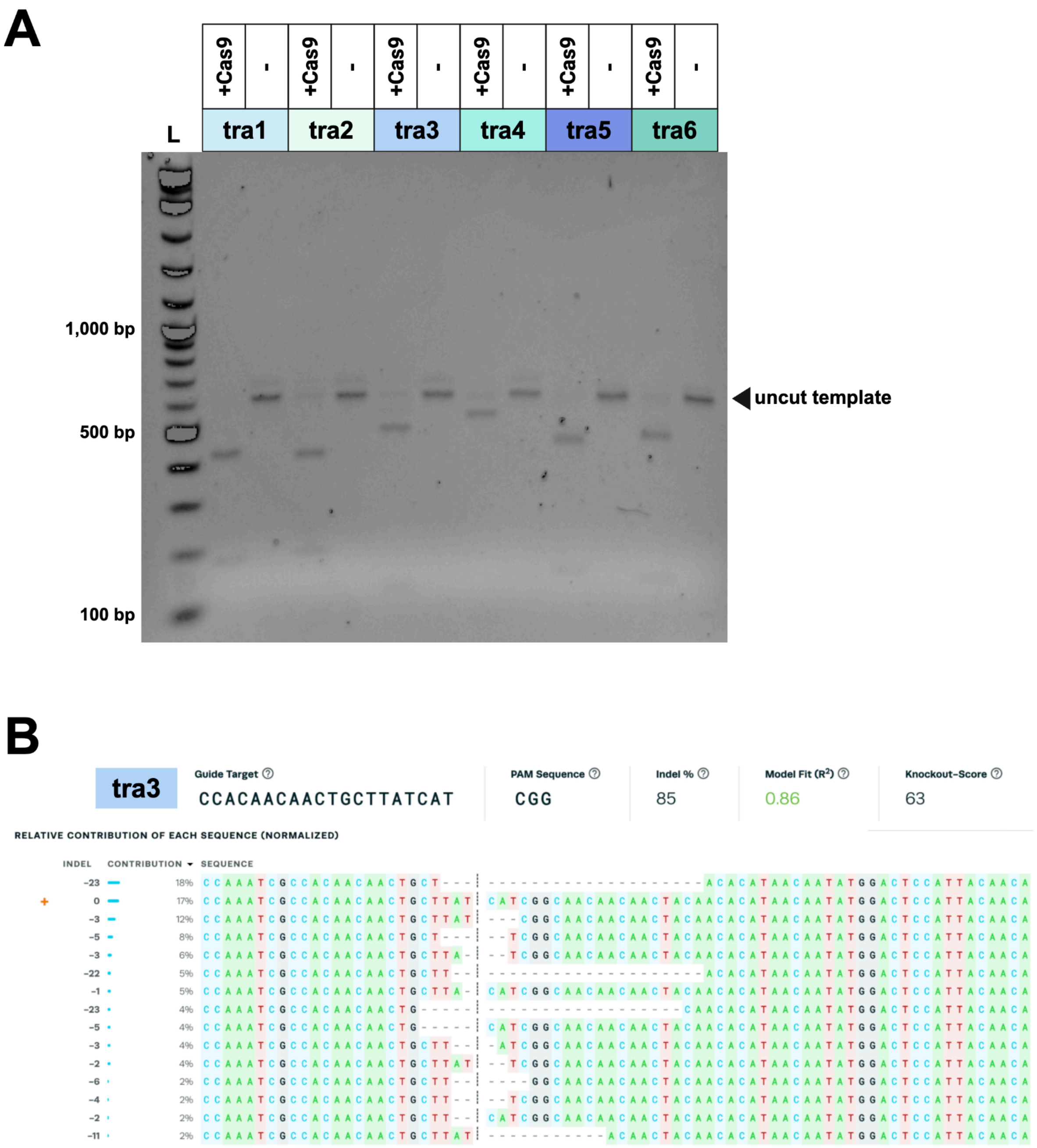


**FIG S1.** Evaluation of sgRNAs targeting *Lctra in vitro* and *in vivo*. **(A)** Gel showing products of *in vitro* Cas9 cleavage assays using sgRNAs *tra1*-*tra6*. Minus signs above lanes indicate negative controls (reaction mixtures without Cas9). **(B)** EditCo ICE analysis of *Lctra* editing rates in wild-type flies that developed from embryos that had been coinjected with *tra3* sgRNA and Cas9 *in vivo*.

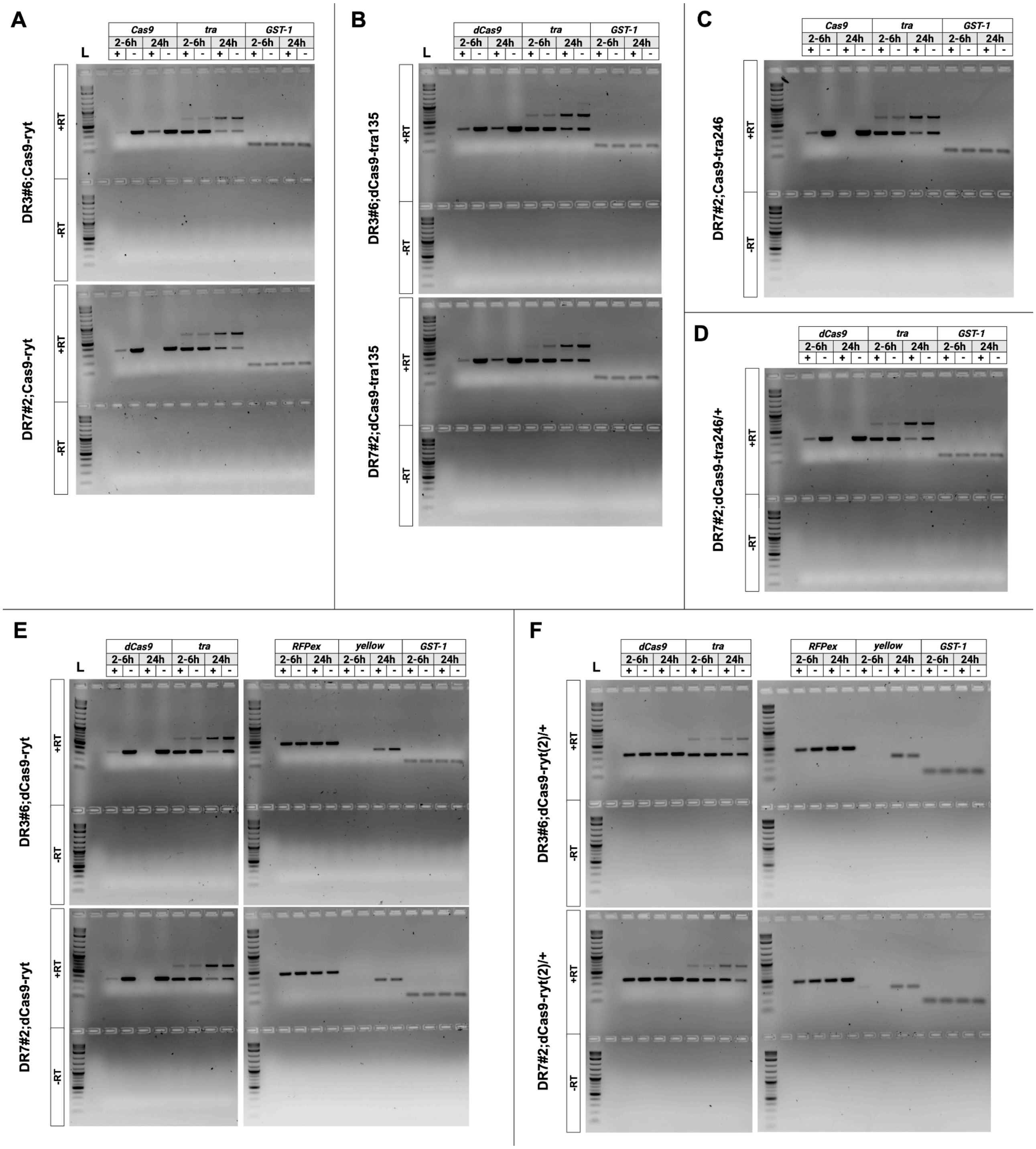


**FIG S2.** Additional gels from RT-PCR of *Cas9/dCas9*, sgRNA arrays, and target genes in sex transformation strains. RT-PCR analysis of **(A-B)** *Cas9* and *Lctra* expression from Cas9 strains and (C-H) *dCas9*, *Lctra*, *RFPex*, and *yellow* expression in dCas9 strains. Blue and pink arrows indicate expected RT-PCR product sizes of male and female splice variants of *transformer*, respectively. The *GST-1* primer pair is used as a positive control. Lanes labeled with + or - denote samples from flies reared on or off 100 µg/mL tetracycline, respectively. 2-6 h = 2-6 h old embryos; 24 h = 24 h old (first instar) larvae. +RT = reactions run with reverse transcriptase; -RT = reactions run without reverse transcriptase (negative controls). L = DNA ladder.

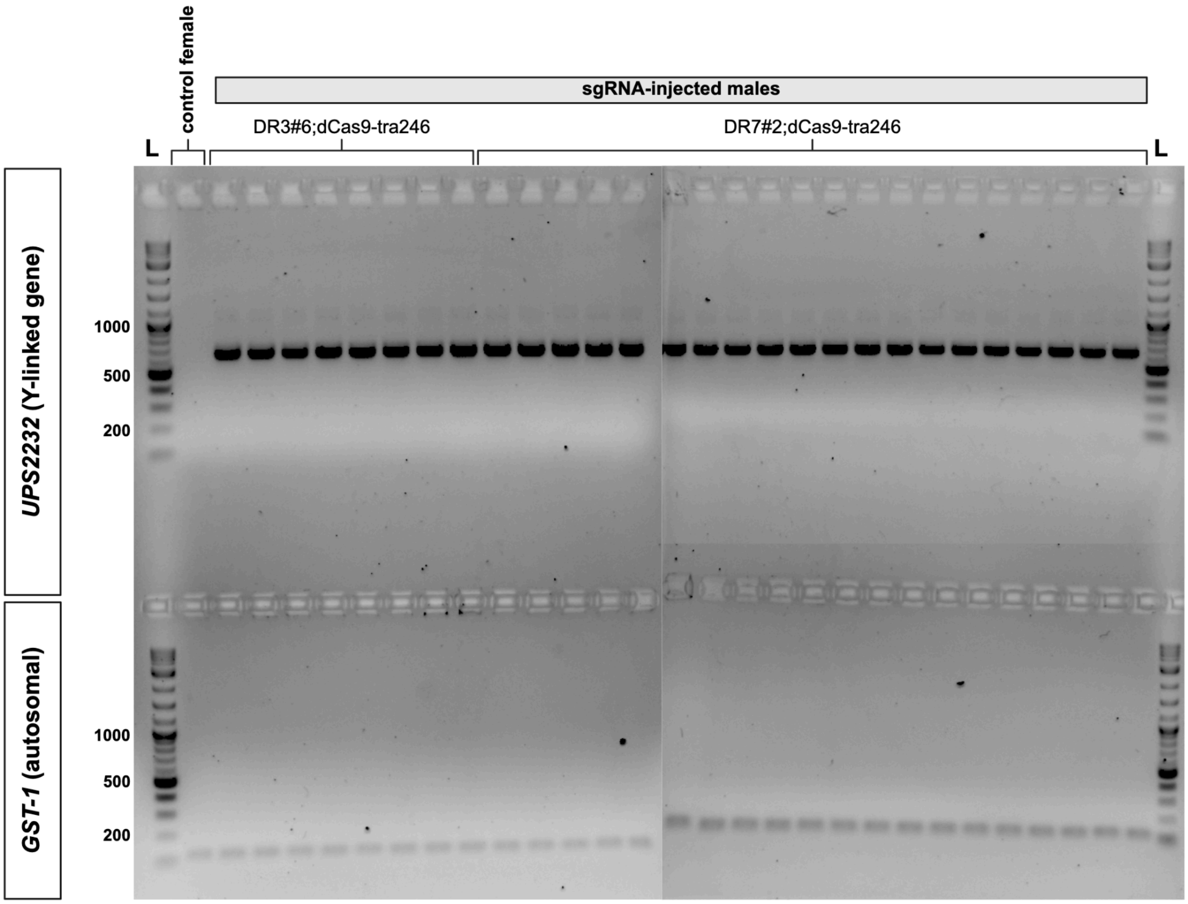


**FIG S3.** Molecular sexing of males from sgRNA-injected dCas9 strains. Top row: Amplification of *UPS2232* (Y-linked gene), which produces a 647-bp product in XY males only. Bottom row: Amplification of *GST-1* gene (autosomal control). L = DNA ladder.
